# Discovery of a *MUC3B* gene reconstructs the membrane mucin gene cluster on human chromosome 7

**DOI:** 10.1101/2021.12.31.474548

**Authors:** Tiange Lang, Thaher Pelaseyed

## Abstract

Human tissue surfaces are coated with mucins, a family of macromolecular sugar-laden proteins serving diverse functions from lubrication to formation of selective biochemical barriers against harmful microorganisms and molecules. Membrane mucins are a distinct group of mucins that are attached to epithelial cell surfaces where they create a dense glycocalyx facing the extracellular environment. All mucin proteins carry long stretches of tandemly repeated sequences that undergo extensive O-linked glycosylation to form linear mucin domains. However, the repetitive nature of mucin domains makes them prone to recombination and render their genetic sequences particularly difficult to read with standard sequencing technologies. As a result, human mucin genes suffer from significant sequence gaps that have hampered investigation of gene function in health and disease. Here we leveraged a recent human genome assembly to identify a previously unmapped *MUC3B* gene located within a cluster of four structurally related membrane mucin genes that we entitle the MUC3 cluster at q22 locus in chromosome 7. We found that *MUC3B* shares high sequence identity with the known *MUC3A* gene, and that the two genes are governed by evolutionarily conserved regulatory elements. Furthermore, we show that *MUC3A, MUC3B, MUC12 and MUC17* in the human MUC3 cluster are exclusively expressed in intestinal epithelial cells. Our results complete existing genetic gaps in the MUC3 cluster that is a conserved genetic unit during primate evolution. We anticipate our results to be the starting point for detection of new polymorphisms in the MUC3 cluster associated with human diseases. Moreover, our study provides the basis for exploration of intestinal mucin gene function in widely used experimental models such as human intestinal organoids and genetic mouse models.

## Introduction

The first draft of the human genome published twenty years ago offered a unique opportunity to decipher the causal relationship between gene sequence, gene function and disease biology (Craig Venter et al., 2001; Lander et al., 2001). But reading and measuring repetitive genomic elements has remained a major technological challenge that has left the human genome riddled with significant sequence gaps. Mucin (*MUC*) genes are characterized by subexonic repeats, characterized by multiple repeated short DNA sequences within coding exons. The resulting tandemly repeated sequences encode long domains rich in proline, threonine and serine (PTS) residues (Lang et al., 2007). Mucin-type tandem repeats undergo O-linked glycosylation on serines and threonines to form densely O-glycosylated linear mucin domains (Nason et al., 2021). The number of tandem repeats and their sequence identity vary between *MUC* genes and are further confounded by considerable length polymorphism between individuals, resulting in variable number tandem repeats (VNTRs). However, VNTRs present a major challenge in analyzing mucin sequences as their repetitive nature and size in several kilobases cause intrinsic instabilities that make them difficult to maintain in bacterial artificial chromosome clones. Consequently, mucin gene VNTRs are underrepresented in human genome assemblies (Chaisson et al., 2015) and continue to hamper efforts to investigation of *MUC* gene function.

Mucins are an ancient family of proteins in the animal kingdom. The earliest mucin genes appeared 700-800 million years ago in primitive marine metazoans such as sea anemones, sponges and jelly combs and have since expanded to all branches of the tree of life (Hedges et al., 2006; Lang et al., 2016). Currently, the human mucin family consists of secreted gel-forming mucins (MUC2, MUC5A, MUC5B, MUC5AC, MUC6, MUC7 and MUC20) and a distinct subfamily of membrane mucins (MUC1, MUC3, MUC4, MUC12, MUC13, MUC15, MUC16, MUC17, MUC21 and MUC22) that are inserted into cell membranes via a transmembrane domain (Pelaseyed and Hansson, 2020). Membrane mucins are single-pass type I transmembrane proteins that are guided to the secretory pathway via an N-terminal signal sequence. In the endoplasmic reticulum, membrane mucins undergoes N-linked glycosylation and folding, which in most cases requires a strain-dependent autocatalytic cleavage at a Sea urchin sperm protein, Enterokinase and Agrin (SEA) domain (Ligtenberg et al., 1992). The cleaved protein fragments remains non-covalently attached at the SEA domain as the mucin protein transits to the Golgi apparatus for O-linked glycosylation. Consequently, the mature SEA-type membrane mucin reach the plasma membrane as heterodimer of a glycosylated extracellular N-terminal subunit that remains non-covalently linked to a membrane-attached C-terminal subunit. The evolutionary origins of the SEA domain date back to single-celled eukaryotes while SEA-type membrane mucins emerged in vertebrates (Lang et al., 2007; Pei and Grishin, 2017). While our earlier work has shown that the SEA domain undergoes conformational unfolding in response to mechanical tension, its ultimate function remains elusive (Pelaseyed et al., 2013).

In humans, membrane mucins genes *MUC3, MUC12* and *MUC17* map to the chromosomal locus 7q22. The three genes are arranged in a *MUC3*-*MUC12*-*MUC17* cluster (hereafter called MUC3 cluster), and flanked by *ACHE* upstream of *MUC3*, and *TRIM56 and SERPINE1* downstream of *MUC17*. The stereotypic *ACHE-MUC3*-*MUC12*-*MUC17*-*TRIM56-SERPINE1* unit is highly conserved in vertebrates. *Mus musculus* carries a MUC3 cluster on chromosome 5 where three membrane mucin genes are flanked by *Ache* and *Trim56-Serpine1*. Notably, the *Muc3* gene in *M. musculus* maps directly upstream of *Trim56* and shares 43% sequence identity with human *MUC17*, but only 28% identity with human *MUC3*, indicating that murine *Muc3* is a homolog of human *MUC17* while the murine homologues for *MUC3* and *MUC12* are poorly defined. The nonmammalian vertebrate *Xenopus tropicalis* carries seven genes encoding SEA-type membrane mucins, of which three are arranged in tandem on chromosome 3 followed by a homolog of human *SERPINE1*, suggesting that the MUC3 cluster first emerged in amphibians (Lang et al., 2007).

The current human genome assembly (GRCh38.p13) is estimated to contain unsolved gaps corresponding to nearly 150 million base pairs (Mbp) (Nurk et al., 2021), which we postulate underlie the lack of complete sequences for human *MUC* genes in general and the MUC3 cluster in particular. In this work, we take advantage of the most recent T2T-CHM13 assembly of the human genome (Nurk et al., 2021) to provide evidence for the existence of a *MUC3B* genes in the human genome. We also demonstrate that *MUC3A* and *MUC3B* genes are conserved in late hominoids such as the chimpanzee as well as Old World monkeys. Finally, by exploring published RNA-sequencing data sets, and applying quantitative gene expression analysis in human tissues, we show that *MUC3A* and *MUC3B* expression is limited to intestinal epithelial cells.

## Results

### The evolution of a MUC3 cluster in Cercopithecoids and Hominoids

The human chromosome locus 7q22 contains three *MUC* genes *MUC3, MUC12* and *MUC17*, arranged in a MUC3 cluster flanked by *ACHE* at its 5’ end, and *TRIM56* and *SERPINE1* at its 3’ end (Figure 1A). Using *ACHE, TRIM56* and *SERPINE1* as genomic markers, we identified the MUC3 cluster in species belonging to the Catarrhini parvorder, namely cercopithecoid (Old World monkeys) and hominoid superfamilies, the latter including the genera *Pongo (*orangutang*), Gorilla, Pan* (chimpanzee and bonobo) and *Homo* (Figure 1B). In cercopithecoids we identified a MUC3 cluster with a length of 153 kilo base pairs (kbp) in rhesus (*Macaca mulatta*), while the corresponding gene cluster in the baboon (*Papio Anubis)* consisted of two mapped sequences with a total length of 138 kbp (Figure 1B). In hominoids, MUC3 cluster length ranged from 106 kbp in the gibbon (*Nomascus leucogenys*) to 203 kbp in the orangutang (*Pongo abelii*). In the Homininae subfamily, we observed striking differences between the MUC3 cluster length in chimpanzee (*Pan troglodytes*) and its two closest relatives; the gene cluster in *H. sapiens* GRCh38 assembly was 73 kbp shorter and, in the gorilla (*G. gorilla*) 66 kbp shorter, than in the chimpanzee (Figure 1B). Thus, we hypothesized that the current human GRCh38.p13 assembly contains significant gaps within the MUC3 cluster.

**Figure 1.**
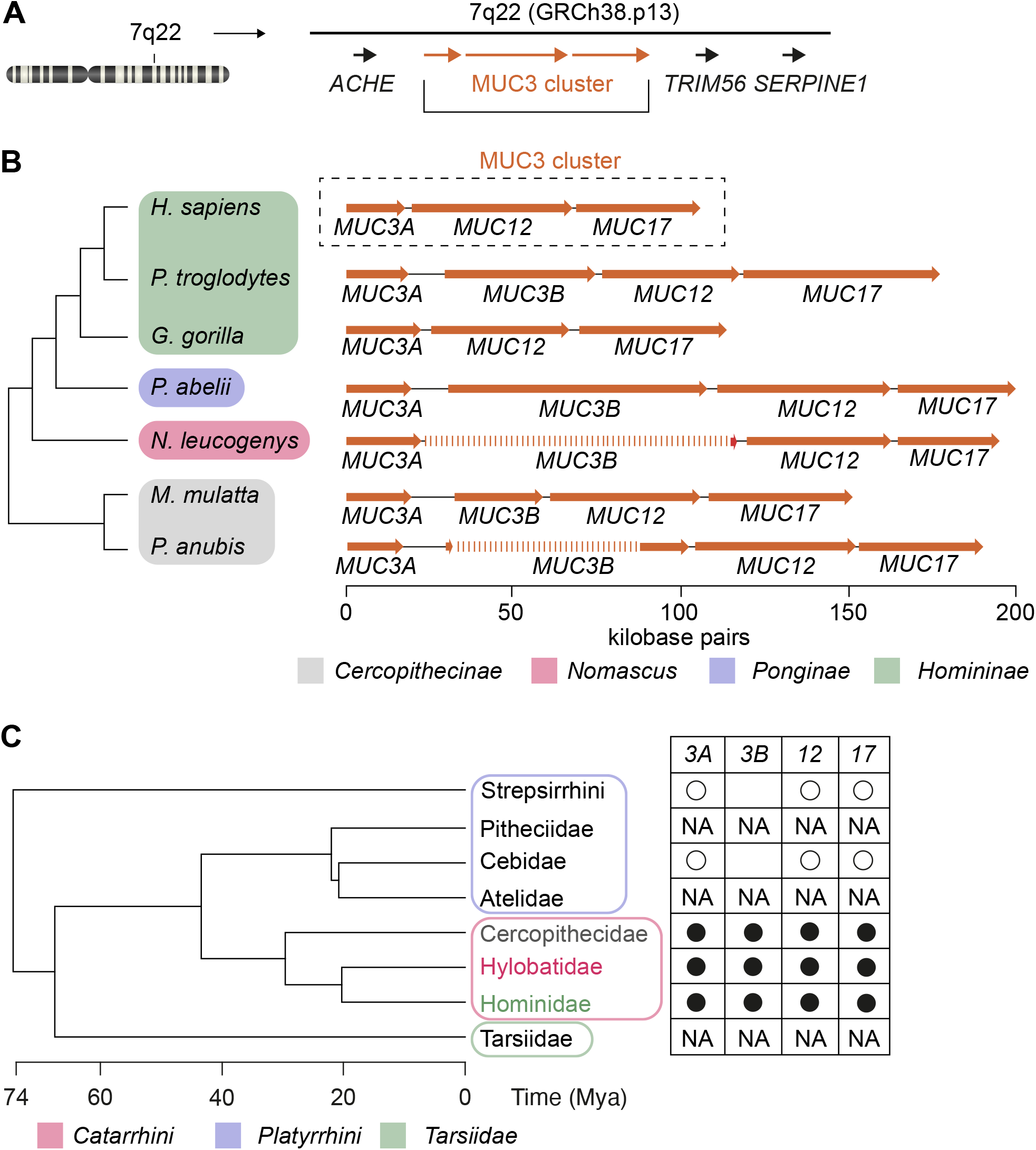
Conservation of MUC3 cluster in Cercopithecoid and Hominoid superfamilies. A) MUC3 cluster in the current GRCh38.p12 assembly is flanked by genes ACHE, TRIM56 and SERPINE1 at locus q22 in human chromosome 7. B) Members of the cercopithecoid and hominoid superfamilies, with the exception of *H. sapiens* and *G. gorilla*, carry a MUC3 cluster consisting of *MUC3A, MUC3B, MUC12* and *MUC17* genes. C) Emergence of *MUC3B* gene in MUC3 cluster in Catarrhini parvorder (filled black circles). Open circles indicated presence of *MUC3, MUC12* and *MUC17* genes in Platyrrhini parvorder. NA indicates lack of sufficient sequence information for detection of MUC3 cluster.

To test our hypothesis, we used a set of defined criteria for exploring distinct *MUC* genes within MUC3 clusters in available primate genome assemblies. We scanned the clusters for 1) start codons, 2) long mucin-type PTS-encoding exons, 3) conserved SEA domains specific for membrane mucins and, 4) unique intronic and exonic sequences that separate individual *MUC* genes. Our analysis revealed that all cercopithecoids carried a MUC3 cluster consisting of *MUC3, MUC12* and *MUC17* genes (Figure 1B). Strikingly, the primate *MUC3* gene existed as two distinct *MUC3A* and *MUC3B* genes, although only partial sequences of the N- and C-terminal fragments of a *MUC3B* gene were identified in *P. anubis*. The hominoid superfamily, with the exception of *H. sapiens* and *G. gorilla*, carried a *MUC3A* gene and full or partial sequences of *MUC3B*. By exploring all existing primate sequences flanked by *ACHE, TRIM56* and *SERPINE1*, we concluded that a *MUC3B* gene emerged exclusively in the Catarrhini parvorder 29.4 million years ago (Mya) (Figure 1C).

### The human 7q22 locus contains a *MUC3B* gene

As humans and chimpanzees share 98.8% of their genomic DNA and the chimpanzee genome carries a *MUC3B* gene in the MUC3 cluster, we hypothesized that absence of a *MUC3B* gene at the human 7q22 locus is a result of sequence gaps in the publicly available genome assembly. In the quest of a human *MUC3B* gene, we explored PacBio Single Molecule Real-Time (SMRT) reads from a human HX1 (Shi et al., 2016) and identified 3 individual reads that covered the 3’ end region of *MUC3A (*encoding the C-terminal region of MUC3A protein, designated *MUC3A* C-term), an intergenic region and the 5’ end region of a putative *MUC3B* gene (designated *MUC3B* N-term) (Figure 2A). Strikingly, the length of intergenic region was in average 10,939 bp, which corresponded to the length of *MUC3A*-*MUC3B* intergenic region in catarrhines (average 11,810 bp). Moreover, we identified 5 SMRT reads covering *MUC3B* C-term, an intergenic region and *MUC12* N-term (Figure 2A). The average length of *MUC3B-MUC12* intergenic region was 2469 bp and conserved in catarrhines (average 2491 bp). This initial exploration provided evidence for the existence of a distinct human *MUC3B* gene. However, because *MUC3A* and *MUC3B* share high sequence identity (87% and 94% for N-term and C-term across catarrhines) and the error rate of the SMRT reads was 70-85%, the HX1 assembly could not with high confidence distinguish between the two *MUC3* genes. Moreover, the reads failed to capture the length and sequence identity of a predicted single PTS-encoding exon in *MUC3B*.

**Figure 2.**
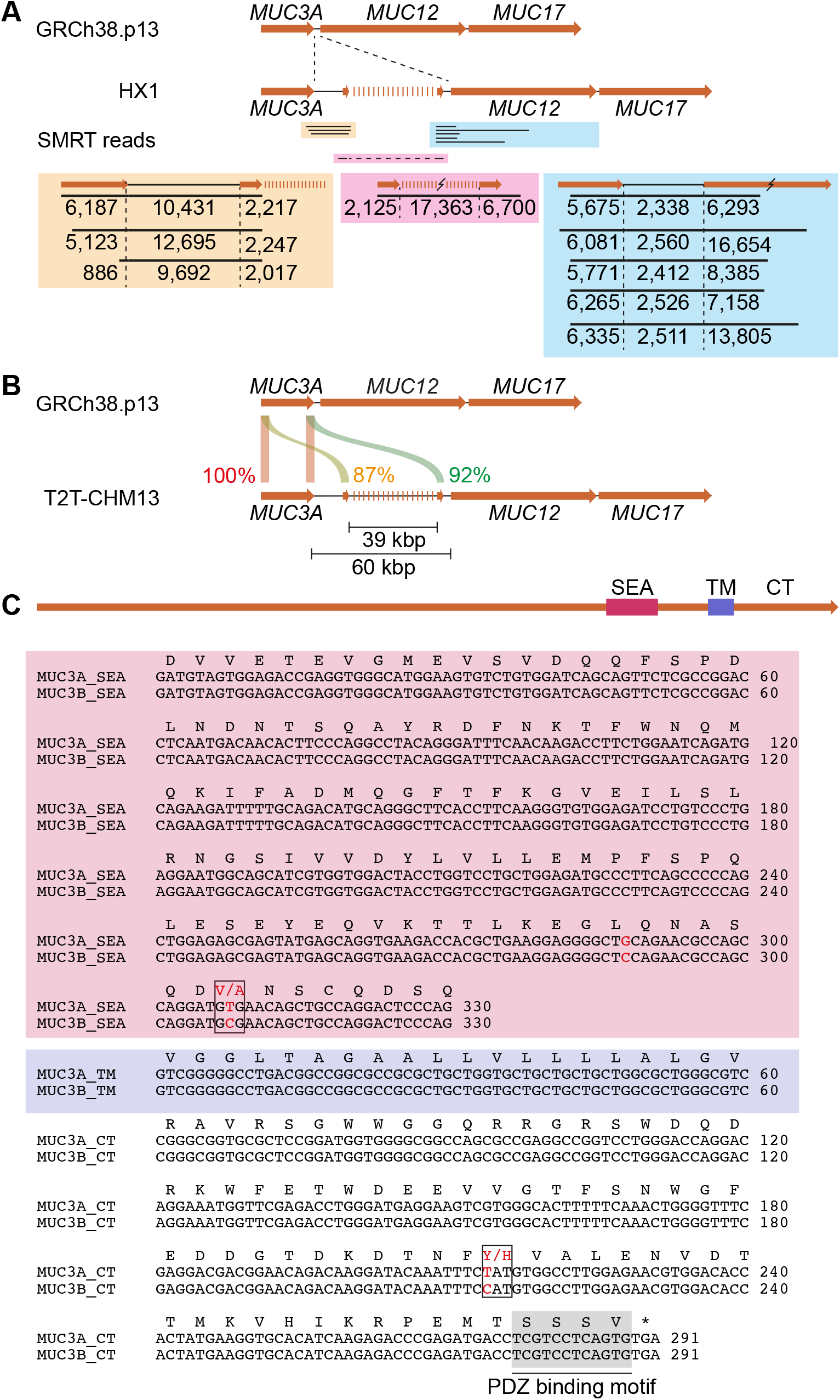
Evidence of a putative *MUC3B* gene in recent human genome assemblies. A) Exploration of PacBio sequencing of HX1 genome identified SMRT reads covering the N-intergenic sequences between *MUC3A* and putative *MUC3B*, an incomplete PTS sequence and intergenic sequences between putative MUC3B and MUC12. B) The T22-CHM13 assembly contains a 60 kb gap between MUC3A and MUC12. C) Sequence alignment of SEA, transmembrane (TM) and cytoplasmic tails (CT) of *MUC3A* and putative *MUC3B* shows high sequence identity, nucleotide mismatches and a conserved PDZ binding motif.

The current GRCh38.p13 draft covers lightly packed euchromatic regions corresponding to 92% of the human genome, while more complex regions including long tandems repeats in *MUC* genes are underrepresented. A recently published CHM13 T2T v1.1 assembly, based on long-read genome sequencing of homozygous complete hydatidiform mole (CHM) cells followed by gapless telomere-to-telomere assembly, adds approximately 200 Mbp to the GRCh38.p13 assembly (Nurk et al., 2021). Importantly, the T2T-CHM13 assembly revealed a 60 kbp gap between *MUC3A* and *MUC12* at locus 7q22 (Figure 2B). Within this gap, we identified a 39,267 bp long PTS-encoding exon flanked upstream by a 2,187 bp long sequence with 87% identity to *MUC3A* N-term. Downstream of the PTS-encoding exon, we identified a 6,303 bp long sequence that was 92% identical with *MUC3A* C-term, and contained a SEA domain, a transmembrane domain and a cytoplasmic tail with a conserved PDZ motif (Malmberg et al., 2008) (Figure 2C). This data suggests that the T2T-CHM13 assembly contains a putative *MUC3B* gene at locus 7q22 with high sequence identity with the annotated *MUC3A* gene.

### Distinct human *MUC3A* and *MUC3B* genes share high sequence homology

To better characterize a putative human *MUC3B* gene, we compared the exon-intron structure of the new *MUC3B* gene to *MUC3A*. A previous study reported 11 exons in a putative *MUC3B* gene (Pratt et al., 2000). Our analysis of T2T-CHM13 assembly revealed that *MUC3A* and *MUC3B* both have 12 exons with near identical nucleotide lengths, with the exception of the PTS-encoding exon 2 that is 15,873 bp (5,291 amino acids) in *MUC3A* and 39,267 bp (13,089 amino acids) in *MUC3B* (Figure 3A). Nucleotide sequence identity amongst individual exons of *MUC3A* and *MUC3B* was in average 93%, and 92% for introns. The superfamilies of hominoids (apes and humans) and cercopithecoids (Old World monkeys) diverged around 29 Mya (Hedges et al., 2006). Sequence alignments between N-terminal and C-terminal of *MUC* genes in the MUC3 cluster showed a high degree of conservation between *H. sapiens* and members of the cercopithecoid and hominoid branches. Human *MUC3A* N-term was 99% identical to chimpanzee *MUC3A* N-term and 90-91% identical to *MUC3A* N-term in the cercopithecoid members rhesus and baboon. *MUC3A* C-term showed a slightly higher degree of divergence compared to MUC3A N-term (Figure 3B, Supplementary table 1). The same trend was observed for *MUC3B*, in which the *MUC3B* C-term was less conserved that *MUC3B* N-term within the hominoid and cercopithecoid superfamilies. Tandem repeat regions are prone to duplications and deletions caused by recombination (Svensson et al., 2018). Accordingly, we observed higher evolutionary sequence divergence in the PTS-encoding exon 2 of *MUC* genes in the MUC3 cluster (Figure 3B, Supplementary table 1). Moreover, pairwise alignment of Catarrhini MUC3 cluster genes revealed a general trend towards expansion of tandem repeats during primate evolution (Figure S1). Specifically within exon 2 of human *MUC3A* and *MUC3B*, we identified imperfect repeats with 87% amino acid sequence identity between *MUC3A* and *MUC3B*. In addition, *MUC3B* contained an additional 1368 amino acids of imperfect repeats (Figure 3C). *MUC3A* and *MUC3B* also harbored 166 and 549 perfect tandem repeats of a 17 amino acids long consensus sequence (ITTTETTSHSTPSFTSS) (Figure 3D). Thus, we conclude that the genetic structure of *MUC3B* is highly similar to *MUC3A* and that the two *MUC3* genes are likely paralogous genes characterized by variable number of tandem repeats.

**Figure 3.**
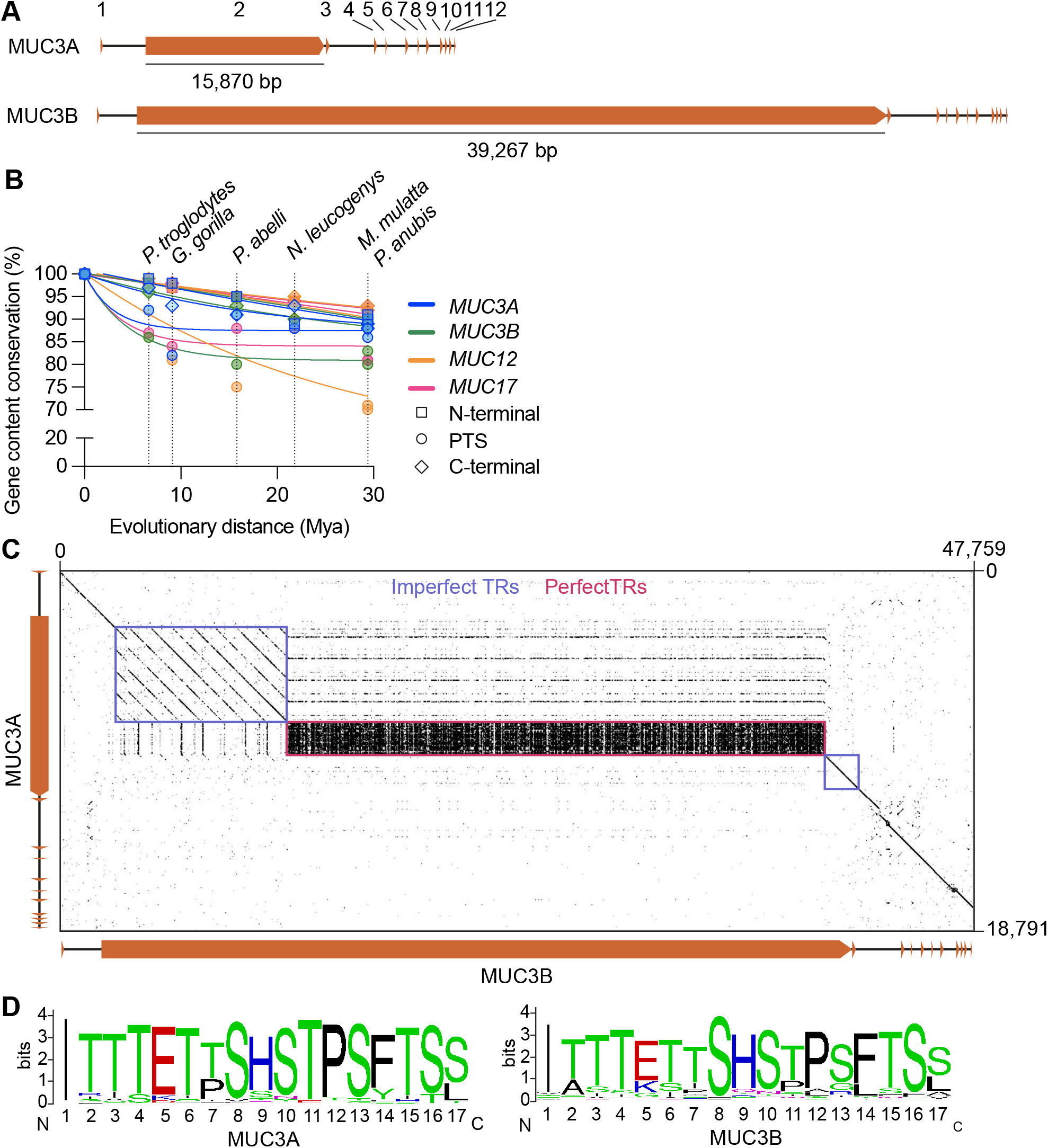
Comparison of genetic and structural features of *MUC3A* and *MUC3B* genes. A) Exon structure and length of exon 2 of *MUC3A* and *MUC3B*. B) Evolutionary rate of N-terminal-, PTS- and C-terminal-encoding exons in *MUC3A, MUC3B, MUC12* and *MUC17* measured as gene content conservation (%) versus evolutionary distance (Mya). C) Dotplot of pairwise sequence alignment of *MUC3A* and *MUC3B* identified imperfect (blue) and perfect (red) tandem repeat sequences in exon 2. D) Sequence logo representation of 166 and 549 perfect tandem repeats (TRs) in exon 2 of *MUC3A* and *MUC3B*, respectively.

### *MUC3A* and *MUC3B* are regulated by conserved regulatory elements

Sequences upstream of a gene’s transcription start site (TSS) contain regulatory elements that dictate gene expression, yet the regulatory elements for *MUC3A* and *MUC3B* genes have not been characterized. Sequence analysis of presumed regulatory sequences -1 kbp of *MUC3A* TSS identified a candidate cis-Regulatory Element (cCRE) at position -403 bp (Figure 4A) that shared 83% identity with the corresponding region in *MUC3B*. Published DNase I hypersensitive site sequencing (DNase-seq) data sets from human small intestine and colon revealed high DNase hypersensitivity signals within the cCRE (Figure 4A). Moreover, we identified high signals for active chromatin markers H3K9ac and H3K4me3 within the *MUC3A* cCRE in human small intestine and colon, while active chromatin signals in the stomach were either not detected or low. By predicting transcription factor binding sites (TFBSs) using the JASPAR CORE vertebrate collection (Fornes et al., 2020; Sandelin et al., 2004), we identified putative TFBSs in cCRE of *MUC3A* (Figure S2). Seven of these transcription factors (ELF3, HNF4A, HNF4G, KLF4, PPARA, STAT3 and XBP1) are enriched in transporting intestinal epithelial cells in human and mouse intestine (Figure 4B, Figure S3). An alignment of putative promoter regions upstream of *MUC3A* and *MUC3B* genes in hominoid and cercopithecoid superfamilies identified conserved TFBSs for HNF4A, HNF4G and STAT3, strongly suggesting that the two *MUC3* genes share an evolutionarily conserved regulatory expression program in the small intestine and colon (Figure 4C, Figure S4, Supplementary table 2).

**Figure 4.**
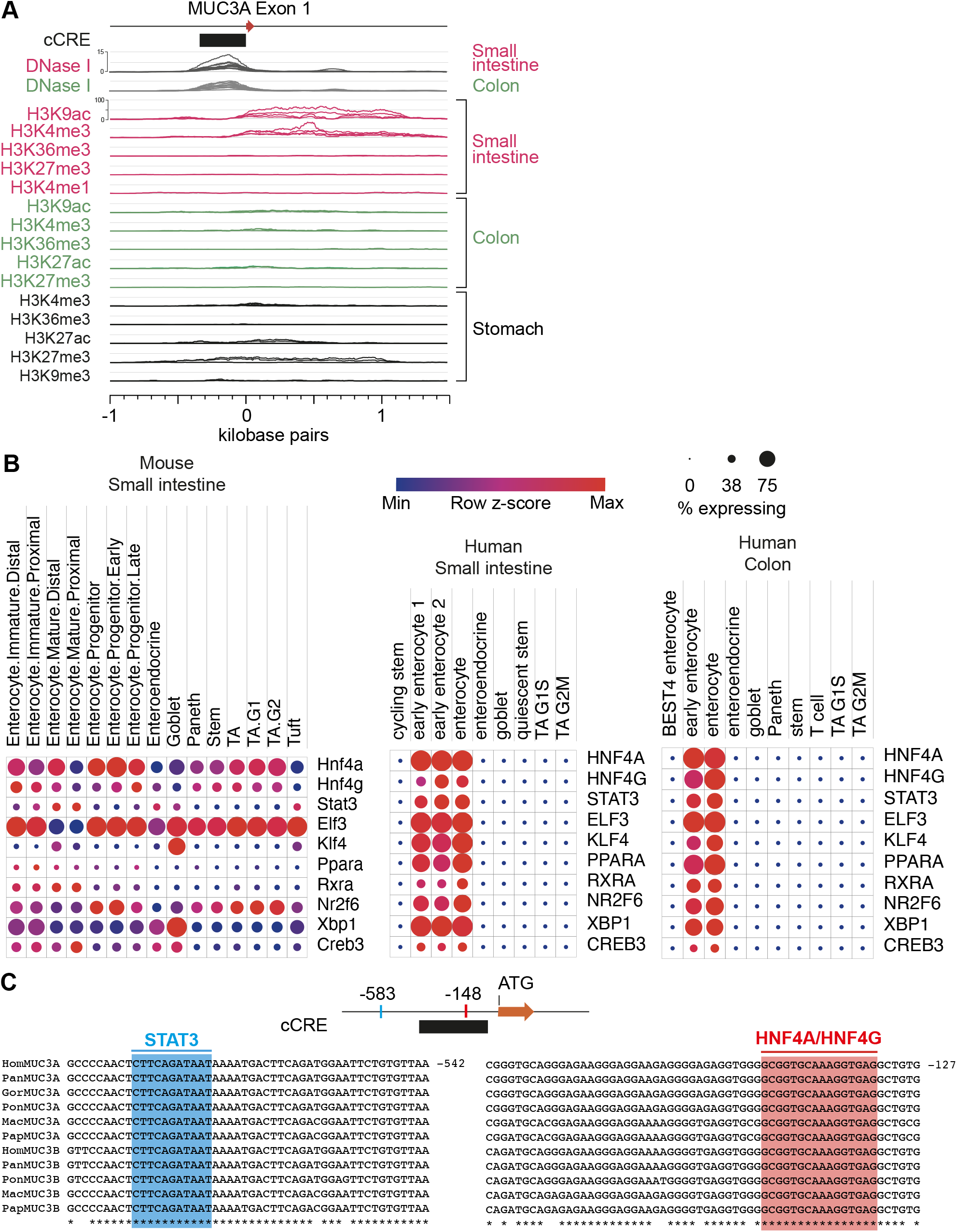
Conserved regulatory elements upstream of *MUC3A* and *MUC3B* genes. A) Epigenetic analysis of human small intestine and colon reveals a DNase I-sensitive candidate cis-regulatory element (cCRE) and specific histone modifications surrounding *MUC3A* transcription start site. B) Single cell analysis of human and mouse intestine shows gene expression of transcription factors in transporting intestinal epithelial cells (enterocytes and colonocytes) with conserved binding sites upstream of *MUC3A* and *MUC3B*. C) Binding sites for transcription factors STAT3 and HNF4A/G are completely conserved upstream of *MUC3A* and *MUC3B* in cercopithecoid and hominoid superfamilies.

### Expression of human *MUC3A* and *MUC3B* genes in the human intestine

To determine whether *MUC3B* is transcribed into messenger RNA, we mapped published RNA-sequencing data sets from human intestine (Wang et al., 2020), liver (Aizarani et al., 2019) and kidney (Su et al., 2021) to the T2T-CHM13 assembly. A considerable number of sequenced reads from *MUC3A* and *MUC3A* transcripts were detected in human ileum, colon and rectum, while liver and kidney were devoid of transcripts from MUC3 cluster genes (Supplementary table 3). Because the exons encoding the N-terminal, PTS and C-terminal regions of *MUC3A* and *MUC3B* share 87%, 83% and 92% identity and PTS-encoding exons are highly repetitive, we predicted a challenge in detecting sufficient unique sequencing information to accurately assign reads to the correct *MUC3* gene. Therefore, we turned our attention to reads that map to the C-terminal region coded by exons 3-12 of *MUC3A* and *MUC3B*, where we identified in average 3.5±1.6 unique reads per kilobase transcript (RPK) of *MUC3A* and 13.0±4.4 unique RPK of *MUC3B* (Figure 5A, Supplementary table 3). We next used unique and shared reads in the C-terminal region to calculate normalized gene expression of *MUC3A, MUC3B, MUC12* and *MUC17* in human intestine. In ileum, *MUC17* showed significantly higher expression than *MUC3A, MUC3B* and *MUC12*, while MUC12 expression showed a trend towards higher expression in rectum compared to ileum. We detected comparable numbers of *MUC3A* and *MUC3B* transcripts in all three intestinal segments (Figure 5B). We next applied a targeted reverse transcriptase quantitative polymerase chain reaction (RT-qPCR) to validate the presence of unique *MUC3A* and *MUC3B* transcripts in ileum collected from five human patients (Supplementary table 4). For this purpose, we designed gene-specific primer pairs targeting sequences in exons 3 and 8 with 9.5-12.0% mismatch between *MUC3A* and *MUC3B*. The resulting 645 bp cDNA amplicons from each transcript were further distinguishable by a unique *PstI* restriction site in *MUC3A* cDNA amplicon (Figure 5C). RT-qPCR from all five patients resulted in the expected 646 bp cDNA amplicon and subsequent *PstI*-digestion produced 380 bp and 266 bp restriction fragments (Figure 5C). Notably, we observed significant differences in intensities of *PstI*-sensitive and *PstI*-resistant fragments produced by the two gene-specific primer pairs. 75% of amplicons generated by *MUC3A*-specific primers were *PstI*-sensitive and therefore originated from *MUC3A* transcripts (Figure 5D). Similarly, 76% of amplicons generated by *MUC3B*-specific primers were *PstI*-resistant *MUC3B* transcripts. Thus, despite significant sequence similarity between the two *MUC3* genes, we successfully identified and distinguished between *MUC3A* and *MUC3B* transcripts in the human intestine.

**Figure 5.**
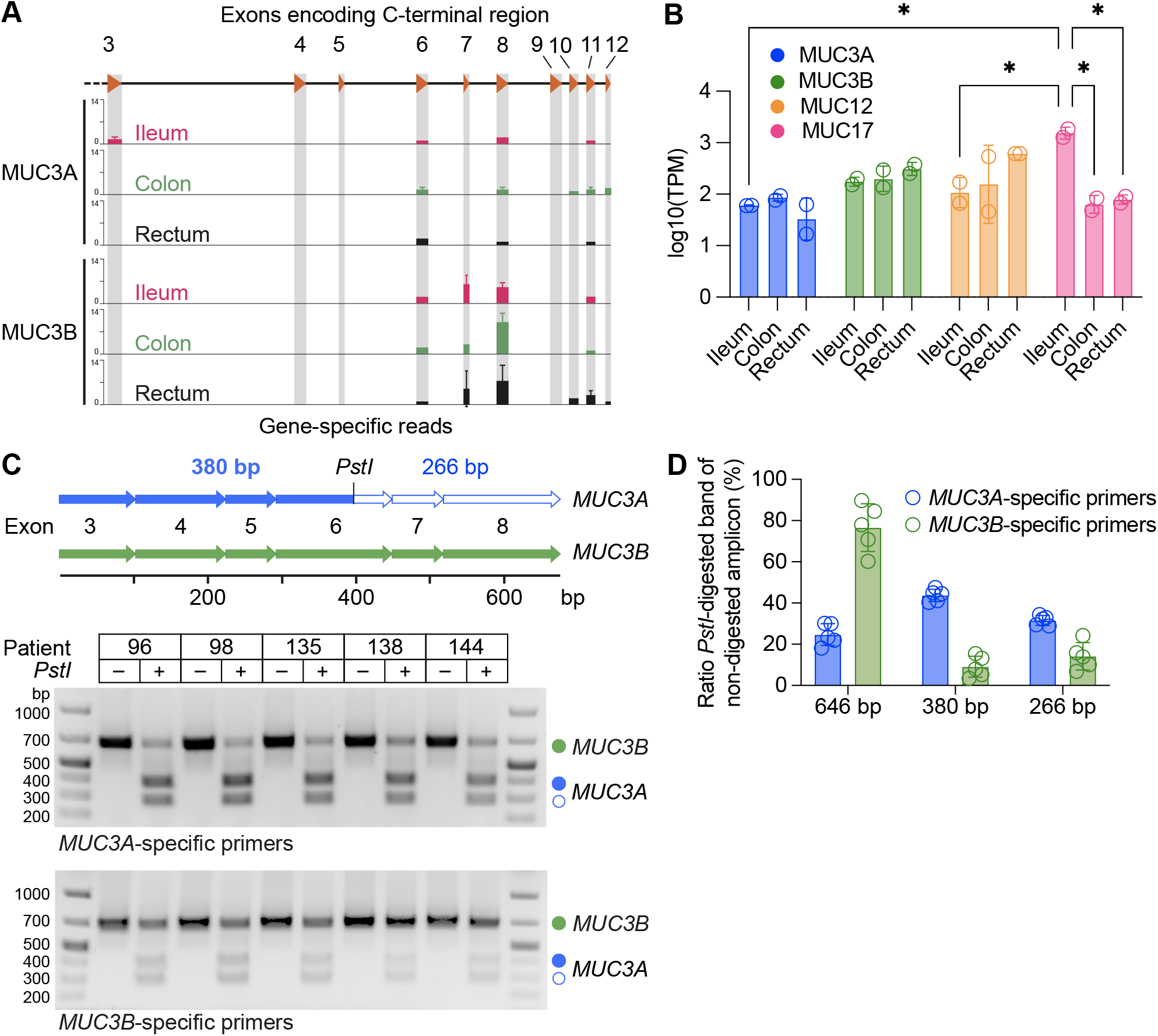
Expression of *MUC3B* gene in human intestine. A) Unique reads for *MUC3A* and *MUC3B* in RNA-sequencing data from human ileum, colon and rectum mapped to T2T-CHM13 human genome assembly. B) Gene expression of MUC3 cluster genes in human ileum, colon and rectum. 2 samples per tissue segments. * p<0.05 as determined by two-way ANOVA, corrected for multiple comparison using Tukey’s test. Data are presented as mean ± standard deviation (SD). C) Specific primers amplify 646 bp cDNAs spanning exons 3-8 in *MUC3A* and *MUC3B* transcripts from ileum of five individuals. *MUC3A* cDNA carries a *PstI* restriction site in exon 6 that distinguishes *MUC3A* from *MUC3B* transcripts. Agarose gel electrophoresis of *PstI* restriction digests of amplified cDNA from *MUC3A* and *MUC3B* transcripts results in 380 bp and 266 bp fragments from *MUC3A* cDNA. D) Quantification of bands from agarose gel in C. n=5 individuals. Data are presented as mean ± standard deviation (SD).

### Completion of a gapless human MUC3 cluster at locus 7q22

Finally, based on the T2T-CHM13 assembly, we revised the gapless length of all four membrane mucins genes in the MUC3 cluster at locus 7q22. In this new assembly, we identified a longer PTS-encoding exon 2 of 15,870 nt in *MUC3A* compared to 8805 nt in GRCh38.p13. The PTS-encoding exon of human MUC12 was 32,428 nt long compared to 14,935 nt in GRCh38.p13 (Supplementary table 5). The complete gapless sequences of *MUC3A, MUC3B, MUC12* and *MUC17* genes at 7q22 locus are publicly available via Mucin database X.X 2021 (http://www.medkem.gu.se/mucinbiology/databases/index.html).

## Discussion

Mucin genes containing long protein-coding sequences consisting of tandem repeats that are difficult to read and measure. As results, many human mucin gene sequences remain uncomplete. Sequence gaps also appear in the genome of *Mus musculus*, an important model organism for understanding human gene function. In an attempt to fill these knowledge gaps in mucin genetics, we focused on a cluster of three membrane mucins genes, the MUC3 cluster, at locus q22 in human chromosome 7. The MUC3 cluster is conserved in the cercopithecoid and hominoid superfamilies, where two distinct *MUC3A* and *MUC3B* genes are annotated in all species except in *H. sapiens* and *G. gorilla*. In this study, we leveraged the recent T2T-CHM13 assembly to fill a 60 kb sequence gap sandwiched between *MUC3A* and *MUC12* genes. Sequence alignments revealed that this membrane mucin gene shares high structural and sequence similarity with *MUC3A*; it consists of a total 12 exons and carries a PTS-encoding exon 2 encompassing imperfect and perfect tandem repeats that were conserved in *MUC3A*. Moreover, within the gap we identified a SEA domain, a transmembrane domain and a cytoplasmic tail with a Class I PDZ motif that is conserved in all annotated membrane mucins of the MUC3 cluster. Importantly, nucleotide mismatches in introns and exons clearly distinguished this putative *MUC3B* gene from *MUC3A*. Also, lengths of intergenic regions spanning across *MUC3A*, the putative *MUC3B* and *MUC12* corresponded to intergenic lengths observed within the MUC3 cluster of cercopithecoids and hominoids.

The evolutionary conservation of *MUC3A* and *MUC3B* genes further suggests that their regulation is conserved in higher mammals. Comparative sequence alignments and available DNase I- and ChIP-seq data sets uncovered a conserved cis-regulatory element upstream of *MUC3B* that included binding sites for transcription factors HNF4A, HNF4G and STAT3. Notably, HNF4A and HNF4G regulate expression of genes that encode critical brush border proteins in the apical membrane domain of transporting intestinal epithelial cells (IECs) (Chen et al., 2021). STAT3 acts downstream of the heteromeric epithelial cell receptor for the cytokine IL-22 that regulates expression of MUC17, which builds the protective glycocalyx barrier atop transporting IECs (Layunta et al., 2021). Finally, mapping published RNA-seq data sets to the T2T-CHM13 assembly identified unique sequencing reads for *MUC3A* and *MUC3B* genes in the human intestine, while gene expression was absent in liver and kidneys. We also validated high-through put expression data by a targeted quantitative detection of distinct *MUC3A* and *MUC3B* transcripts in the human ileum. Collectively, we identified a previously unannotated *MUC3B* gene at locus 7q22 and provide evidence for its expression in human intestinal epithelial cells.

The *MUC3B* gene emerges first in the parvorder Catarrhini (Old World monkeys) but is absent in Platyrrhini (New World monkeys), suggesting a gene duplication event in the Simian infraorder 43 Mya. Interestingly, N- and C-terminal regions of MUC3A and MUC3B are highly conserved within Catarrhini, whereas the PTS-encoding exons exhibit higher evolutionary divergence. The PTS domains of membrane mucins genes are encoded by short nucleotide sequences organized in tandem repeats. PTS domains are generally poorly conserved (Lang et al., 2007), as individual repeats are added or removed through recombination to generate variable number tandem repeats (VNTRs) that introduce substantial gene polymorphism. In analogy with other genes carrying VNTRs (Sulovari et al., 2019), our study shows that tandem repeat regions of MUC3 cluster genes have undergone expansion during primate evolution. Low conservation and considerable polymorphism between and within species suggest that O-glycosylation of mucin VNTRs is a non-template-driven process that occurs in a specific evolutionary and environmental context with the primary aim of attaching specific glycan epitopes to the mucin protein backbone. Therefore, glycosylation mucin VNTRs in microbe-rich environments such as the oral cavity and gastrointestinal tract are likely under selective pressure to maintain appropriate interactions with microorganisms that have coevolved with the host through various periods of geographical, dietary and life-style adaptations. This co-speciation is exemplified within the human gut microbiome, where the microbiome of present-day humans is enriched in mucin-degrading genes compared to enrichment of starch- and chitin-degrading genes in our ancestral gut microbiome (Wibowo et al., 2021). Another example is found at the epithelial surface glycocalyx, which underwent major remodeling >2 Mya when a human ancestor acquired an inactivating mutation in *CMAH*, a gene responsible for converting N-acetylneuraminic acid (Neu5Ac) to N-glycolylneuraminic acid (Neu5Gc) (Chou et al., 2002). The resulting accumulation of terminal Neu5Ac in the glycocalyx of human cells has since be exploited by numerous pathogenic microorganism such as *Vibrio cholera* (Alisson-Silva et al., 2018).

Due to existing challenges in sequencing very long repetitive regions, the nature of mucin polymorphism and its contribution to human disease phenotypes remains elusive. A recent study showed that the length of VNTRs in membrane mucin MUC1 is associated with several phenotypes related to kidney function (Mukamel et al., 2021), supporting the notion that glycosylated PTS domains of membrane mucins play critical roles in organ function and homeostasis. Intestinal membrane mucin MUC17 is genetically and structurally related with MUC3A and MUC3B and functions as a major building block of the dense glycocalyx covering transporting IECs. In mouse small intestine, Muc17 expression is induced during the suckling-weaning transition when density and complexity of the gut microbiota increases and IECs establish a cell-attached glycocalyx to prevent adhesion of luminal bacteria to the epithelium (Layunta et al., 2021). While the function of the *MUC3A, MUC3B* and *MUC12* remains elusive, their expression varies along different segments of the human intestine, suggesting that MUC3 cluster genes perform segment- and cell-specific functions in humans and other mammalian vertebrates. Our comprehensive map of the MUC3 cluster in the human genome provides opportunities to identify new VNTR polymorphisms associated with disease phenotypes and allow for future exploration of experimental mammalian models such as the mouse for gene orthologs of the MUC3 cluster.

## Methods

### Recruitment of patients and sample collection

Patients ≥18 years who were referred to Sahlgrenska University Hospital (Gothenburg, Sweden) for colonoscopy, were eligible for inclusion and subject to the provision of written informed consent. Patients with macroscopic/microscopic evidence of ileocolonic pathology other than inflammatory bowel disease were excluded. Eight biopsies were obtained from the terminal ileum of each patient. The study protocol was approved by the regional ethics committee (Ethical permit #2020-03196) and was in compliance with the Declaration of Helsinki.

### Phylogenic data

Phylogenic trees and molecular time estimates were extracted from TimeTree (Hedges et al., 2006, 2015).

### Sequence alignments

Local sequence similarity search and identity measurements of *MUC* genes was performed using NCBI BLAST (McGinnis and Madden, 2004). Multiple sequence alignment of *MUC* gene and protein homologues were conducted using CLUSTALW (Larkin et al., 2007). Perl scripts were used for all data extraction (see supplementary information). Promotor regions -1 kb from transcription start site of *MUC3A* and *MUC3B* in cercopithecoid and hominoid superfamilies were aligned using Multiple Alignment using Fast Fourier Transform (MAFFT) high speed multiple sequence alignment tool (Madeira et al., 2019).

### Generation of dotplots for pairwise sequence alignment and sequence logo representations

Dotplots representing pairwise sequence alignments were generated using Genome Pair Rapid Dotter (GEPARD) version 1.40 (Krumsiek et al., 2007). Sequence logos of perfect tandem repeats were generated using WebLogo3 (Crooks et al., 2004).

### Mapping of DNase-seq and ChIP-seq data to the human genome

DNase hypersensitive sequences upstream of *MUC3A* in GRCh38 and Chromatin immunoprecipitation (ChIP) sequencing of human small intestine, colon and stomach samples are summarized in Supplementary table 6 (Zhang et al., 2020). Graphical representation of epigenic signatures was prepared by aggregating multiple segment-sorted tracks using the Matplot function in Washington University Epigenome Browser v53.5.0 (Li et al., 2019).

### Single-cell expression of transcription factors

Expression profiles for transcription factors were extracted from the following data sets available at Single Cell Portal (Broad institute): single-cell transcriptome analysis of human small intestine (GSE148829) (Ziegler et al., 2020), human colon (GSE178341) (Pelka et al., 2021) and mouse small intestine (GSE92332) (Haber et al., 2017).

### Mapping of RNA-sequencing data to T2T-CHM13 human genome assembly

The T2T-CHM13 human genome assembly was downloaded from NCBI BioProject PRJNA559484). Fastq-dump was used to obtain RNA-sequencing reads. Burrows-Wheeler Aligner (BWA) software package (Li and Durbin, 2009) was used to align RNA-sequencing reads to exonic sequences of genes belonging to the MUC3 cluster. Perl scripts were used to perform quality control and measure read number (see supplementary methods). The following publicly available data sets were used to determine *MUC* gene expression in human tissues: single-cell transcriptome analysis of human ileum, colon, rectum (GSE125970) (Wang et al., 2020), human liver (GSE124395) (Aizarani et al., 2019) and human kidney (GSE131685) (Aizarani et al., 2019). Gene expression of individual *MUC* genes in the MUC3 cluster was calculated as transcripts per million (TPM) as previously described (Li et al., 2010).

### RNA extraction from human ileum, cDNA synthesis and RT-qPCR

RNA from human ileal biopsies was extracted using RNeasy Mini Kit (Qiagen). 500 ng of RNA was reverse transcribed to cDNA using TaqMan Reverse Transcription kit (#N8080234, Applied Biosystems) using 2.5 µM random primers using the cycling parameters 25.0°C for 10 min, 37.0°C for 30 min, and 95.0°C for 5 min. 750 ng of cDNA was used for downstream reverse transcription quantitative PCR (RT-qPCR) with 0.3 µM MUC3A-specific primers (forward 5’-TGGGGGTCAGTGGGATGGCCTCAAA-3’; reverse 5’ - CACGTGGGACCGCTCGTCTCC) and MUC3B-specific primers (forward 5’ CGGGGGCCAGTGGGATGGCCTCAAG-3’; reverse 5’-CACGCGGGACCGCTCGTCTCT-3’) using SsoFast EvaGreen Supermix (#1725200, Bio-Rad) on a CFX96 Real-Time PCR Detection System (Bio-Rad) with the cycling parameters 95.0°C for 3 min, 39 cycles of 95.0°C for 10 s, 63.5°C for 10 s, 72.0°C for 20 s. Melting curve analysis was performed at 95.0°C 10 s and 65.0° to 95.0°C at an increment of 5°C for 5 s.

### Restriction site analysis and agarose gel electrophoresis

5 µL of RT-qPCR reaction was digested with 1 µL FastDigest *PstI* restriction enzyme (#FD0614, ThermoFisher Scientific) for 1 h at 37°C. Full-length amplicons and digestion products were separated on 1.5% agarose gel with ethidium bromide.

### Statistics

Statistical analysis and graphical illustrations were performed using GraphPad PRISM 8.3.1 (GraphPad Software). Data are presented as mean ± standard deviation (SD). Statistical tests were applied as indicated. For all statistical analyses: * p<0.05, ** p<0.01, *** p<0.0001, **** p<0.00001, ns = non-significant.

## Author contributions

T.L. collected the data, contributed data and performed analysis. T.P. conceived and designed the analysis, collected data, contributed data, performed analysis and wrote the manuscript.

## Acknowledgements

We thank Professor Gunnar C. Hansson for valuable discussions. This work was supported by the Swedish Society for Medical Research (Svenska Sällskapet för Medicinsk Forskning, grant S17-0005), National Institutes of Health (grants 5U01AI095542-08-WU-19-95 and 5U01AI095542-09-WU-20-77), Wenner-Gren Foundations (grants FT2017-0002, UPD2018-0065, and WUP2017-0005), Jeansson Foundations (grant JS2017-0003), and the Åke Wiberg Foundation (grant M17-0062).

