## Supplementary information for "Discovery of a *MUC3B* gene reconstructs the membrane mucin gene cluster on human chromosome 7"

**Figure S1**

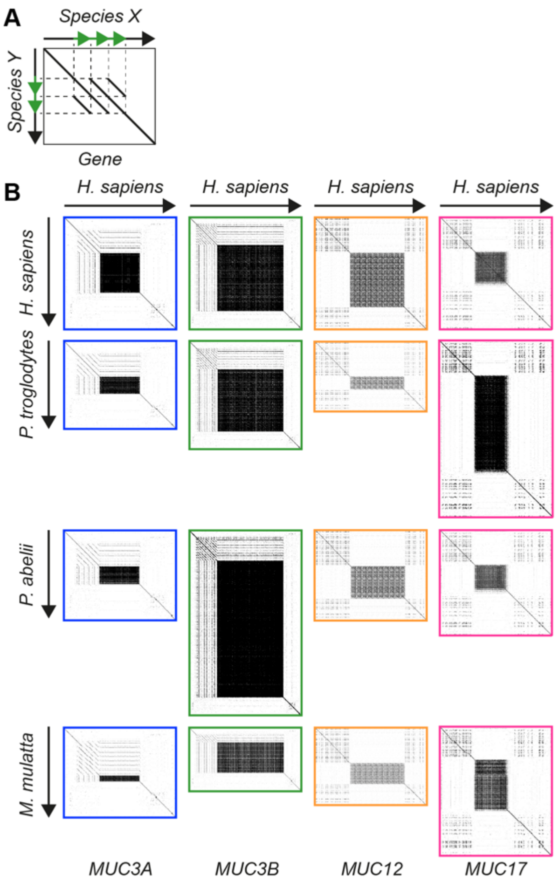


**Supplementary figure S1 related to Figure 3C-D. Dotplots representing pairwise sequence alignments of MUC3 cluster genes belonging to members of Catarrhini parvorder.**

A) Schematic dotplot depicting pairwise sequence alignment of a gene from two distinct species. Tandem repeat structures are characterized by parallel lines. Differences in tandem repeat number result in asymmetric clusters of parallel lines.

B) Dotplot showing pairwise alignment of intronic and exonic sequences in MUC3 cluster genes belonging to *H. sapiens* and three *Homininae*, *Ponginae* and *Cercopithecinae* subfamily species.

**Figure S2**

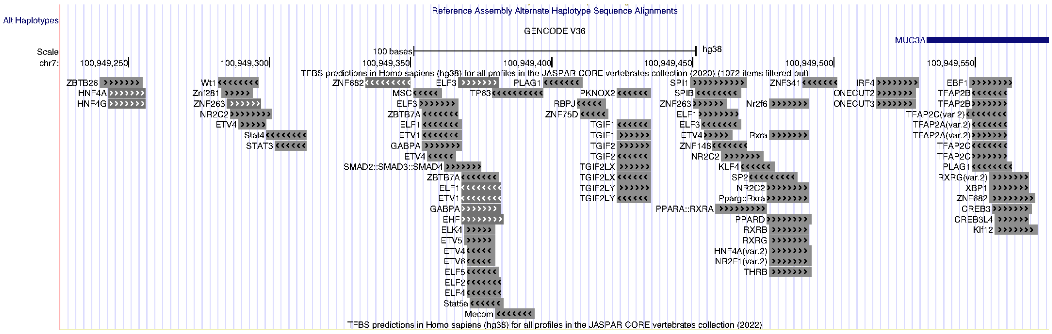


**Supplementary figure S2 related to Figure 4B. Position of transcription factor binding sites upstream of *MUC3A* transcription start site.**

### Figure S3

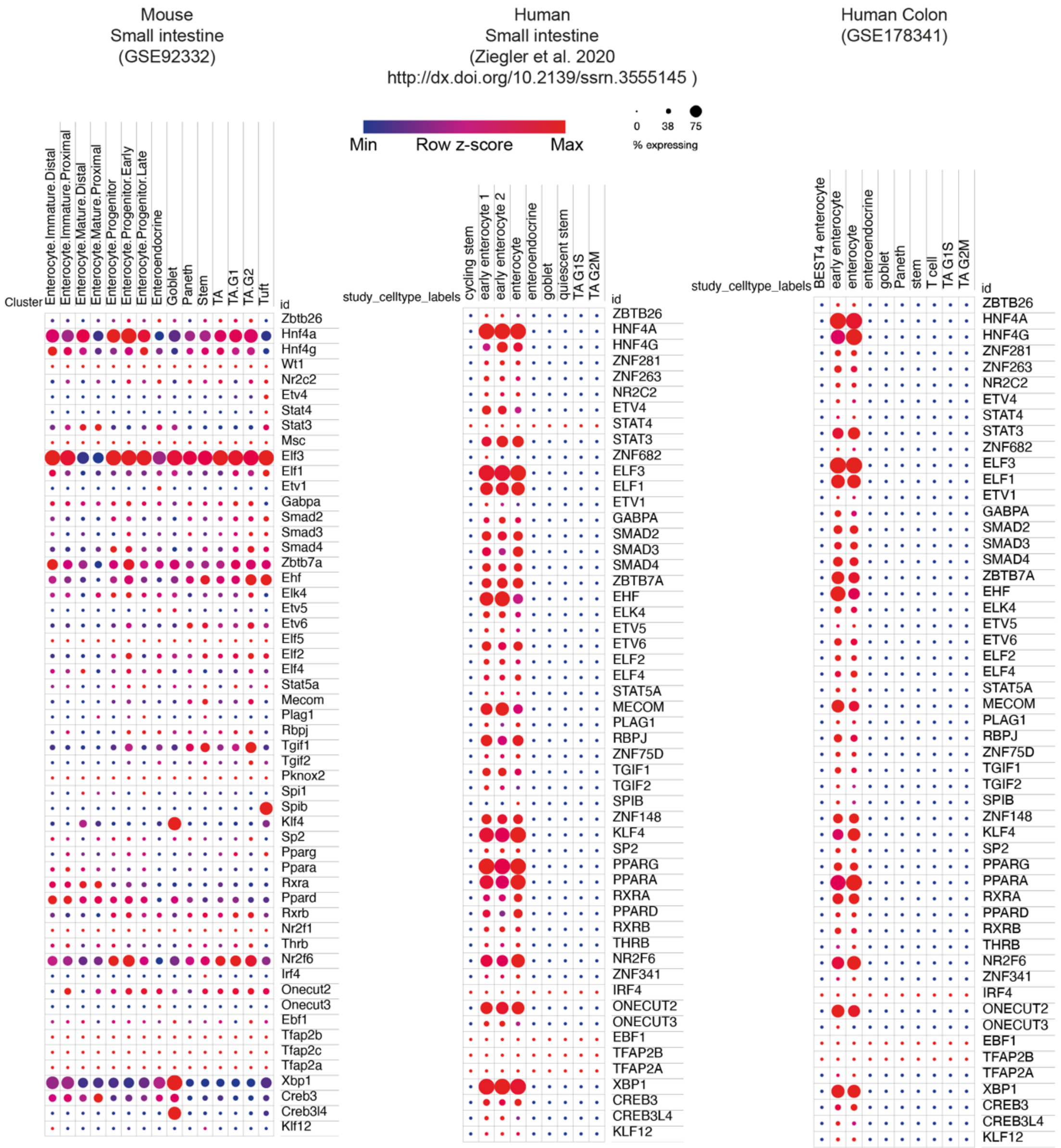


**Supplementary figure S3 related to Figure 4B. Transcription factor expression in human and mouse intestine.**

Summary of single cell gene expression of transcription factors with putative bindings sites upstream of *MUC3A* and *MUC3B* in human small intestine and colon, and mouse small intestine.

**Figure S4**

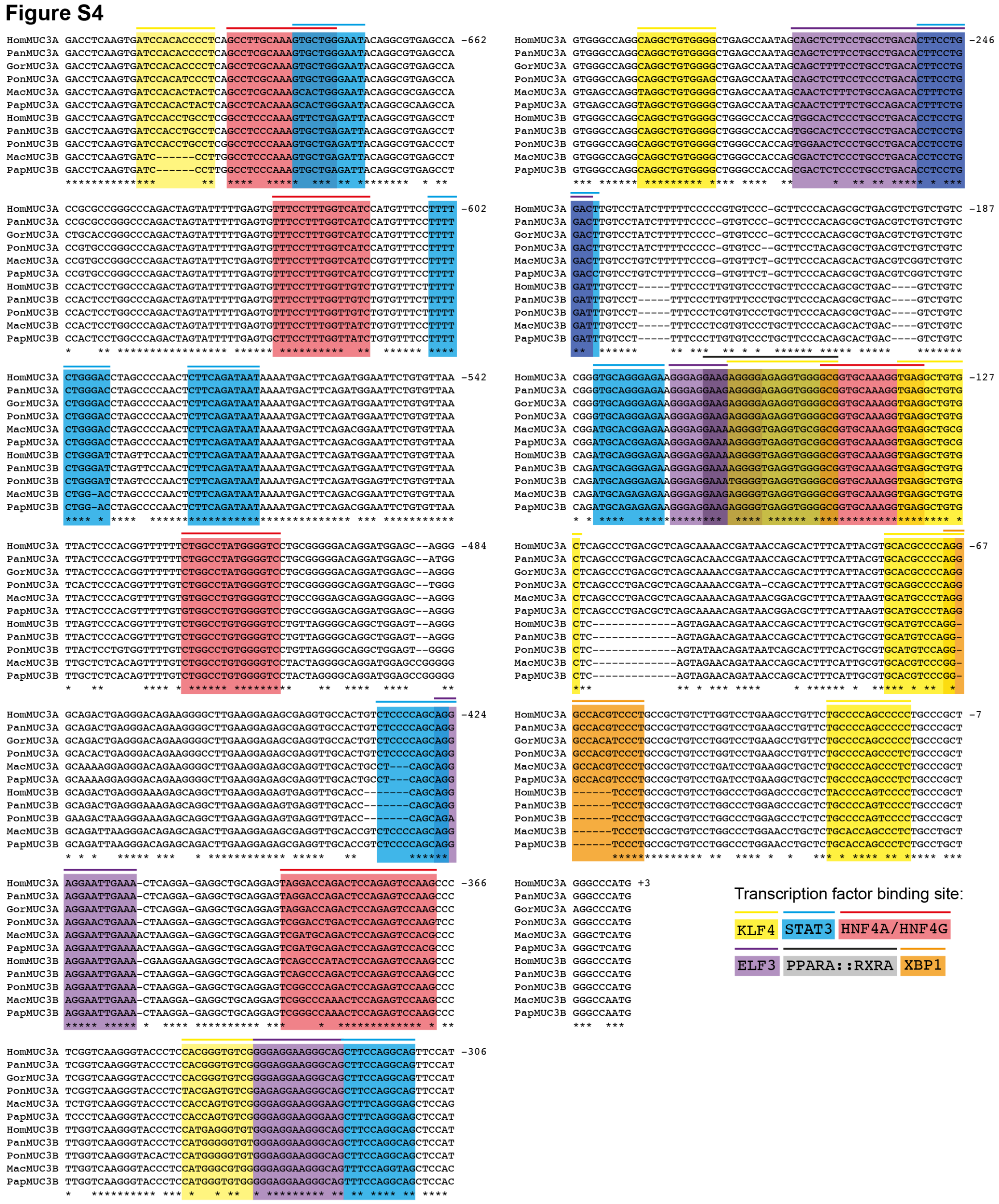


**Supplementary figure S4 related to Figure 4C. Conserved transcription factor binding sites upstream of *MUC3A* and *MUC3B* genes.**

Sequence alignment of sequences upstream of *MUC3A* and *MUC3B* shows conservation of binding sites from seven transcription factors expressed in human enterocytes and colonocytes.

### Supplementary methods

#### Perl scripts for extracting specific sequences based on BLAST results

Perl Script for extraction of fragments using start and end positions obtained by BLAST results

```
#!/usr/bin/env perl
my $begin = $ARGV[0];
my $end = $ARGV[1];
my $source = $ARGV[2];
open (IN, "<$source") ||  die "cannnot open \"$source\": $!";
$OneLineSeq = ();
while (<IN>){
    chomp;
    if (/^>/){print "$_"; print "($begin-$end)\n";}
    else {
        $OneLineSeq = $OneLineSeq.$_;
        $OneLineSeq =~ s/\s//g;
    }
}

$result = substr ($OneLineSeq, $begin-1, $end-$begin+1);
print "$result\n";
close IN;
```

Perl script for extraction of multiple sequences using batch processing based on position file and sequence file

```
#!/usr/bin/env perl

$posfile = shift;
$seqfile = shift;
$OneLineSeq = ();

open SEQFILE, $seqfile or die "Cannot open file\n";
while (<SEQFILE>){
    chomp;
    unless (/>/){
        $OneLineSeq = $OneLineSeq.$_;
        $OneLineSeq =~ s/\s//g;
    }
    else{$id=$_; $id=~s/ .*//;}
}
close SEQFILE;

open POSFILE, $posfile or die "Cannot open file\n";
while (<POSFILE>){
    chomp;
    next if /^\s*$/;
    @position = split (/\.\./,$_);
    $seq = cut ($position[0], $position[1], $OneLineSeq);
    print $id,"(",$position[0],"-",$position[1],")\n",$seq,"\n";
}
close POSFILE;

sub cut {
    my ($begin, $end, $seq)= @_;
    my $result;
    $result = substr ($seq, $begin-1, $end-$begin+1);
    return $result;
}
```

Perl scripts used to perform quality control and measure read number in BWA result files

For single end

```
#!/usr/bin/env perl

while (<>){
    chomp;
    next if /^\@/;
    @line = split (/\t/,$_);
    next if $line[1] >= 256;
    print $_,"\n" if $line[4] >= 20;
}

# Read number could be counted by the number of lines of perl
result file.
```

For paired end

```
#!/usr/bin/env perl

$k=0;
while (<>){
    chomp;
    next if /^\@/;
    @line = split (/\t/,$_);
    $k=$k+1;
    if ($k/2!=int($k/2)){$i=1};
    if ($k/2==int($k/2)){$i=2};
    next if $line[1] >= 256;
    $line=$_;
    $line=~s/\S+\t//;
    print $line[0],"_",$i,"\t",$line,"\n" if $line[4] >= 20;
}

# Read number could be counted by the number of lines of perl
result file.
```

**Supplementary table 1 related to Figure 3B.** Conservation of N-terminal, PTS and C-terminal regions of MUC3 cluster genes in primates.

**Supplementary table 2 related to Figure 4B.** Binding sites for transcription factors enriched in intestinal epithelial cells identified upstream of *MUC3A* gene transcription start site.

**Supplementary table 3 related to Figure 5A-B.** Total reads, unique reads and gene expression of *MUC3A*, *MUC3B*, *MUC12* and *MUC17*, extracted from RNA-sequencing of human ileum, colon, rectum, kidney and liver, mapped to T2T-CHM13.

**Supplementary table 4 related to Figure 5C-D.** Patient demographics.

**Supplementary table 5.** Statistical summary of MUC3 cluster genes belonging to members of Cercopithecoids and Hominoids superfamilies.

**Supplementary table 6 related to Figure 4A.** DNase-sequencing and Chromatin immunoprecipitation (ChIP) sequencing data sets used in this study.
